# Volatile organic compounds of *L. enzymogenes* LEC suppress the growth of fungal and oomycete pathogens and modify the cell physiology of *Rhizoctonia solani*

**DOI:** 10.64898/2026.10.29.762756

**Authors:** Julian Maier, Jannicke Gallinger, Maja Molthan, Natascha Heil, Patrick Schäfer, Ada Linkies

**Affiliations:** Federal Research Centre for Cultivated Plants, Institute for Biological Control, 69221 Dossenheim, Germany; Federal Research Centre for Cultivated Plants, Institute for Plant Protection in Fruit Crops and Viticulture, 69221 Dossenheim, Germany; Geisenheim University, Department of Crop Protection, 65366 Geisenheim, Germany; Justus Liebig University, Institute for Phytopathology, 35392 Giessen, Germany

**Keywords:** phytopathogens, mode of action, biological control, ketones, fluorescence microscopy, transmission electron microscopy

## Abstract

*Lysobacter enzymogenes* LEC is a plant disease-suppressive bacterium. To obtain a more comprehensive understanding of its biocontrol activity and to identify further potential application methods, we investigated the function of its volatile organic compounds (VOCs). VOC activity was assessed *in vitro* using split-plate assays with eight different fungal and oomycete isolates and *in planta* against *Pseudoperonospora cubensis* on cucumber plants. The volatilomes of LEC and the less active *L. enzymogenes* DSM2043 were analysed by GC-MS analyses. Further, the effects of VOCs on cell physiology of *Rhizoctonia solani* were analyzed by fluorescence and transmission electron microscopy. The strongest inhibition of LEC VOCs was observed for *Phytophthora infestans* (94%) and *Phoma lingam* (83%), while inhibitions were lower for *Rhizoctonia solani* (42%) and *Alternaria* spp. (24% and 29%) and not detectable for two *Fusarium* isolates. VOC exposure also significantly reduced downy mildew caused by *P. cubensis* on cucumber plants (39%), although this effect was locally restricted to the leaf directly above the VOC source. In *R. solani*, VOC exposure reduced nuclei abundance and increased lipid accumulation. GC–MS analyses revealed distinct volatile profiles between LEC and *L. enzymogenes* DSM2043 and identified ketones, such as 2-dodecanone and undecanone as candidate bioactive compounds, which is in agreement with previous findings. Our results demonstrate broad-spectrum VOC-mediated biocontrol activity, and thereby provide a more comprehensive understanding of its biocontrol activity and highlight new ways for potential applications of LEC.

**Highlights:**

- Volatile compounds of *L. enzymogenes* LEC inhibit pathogen growth *in vitro*
- Volatile compounds are able to suppress downy mildew in cucumber plants
- The volatilomes of *L. enzymogenes* isolates differ
- Ketones are higher abundant in the volatilome of *L. enzymogenes* LEC
- Volatile compounds modify cell physiology of *R. solani*

## 1. Introduction

Plant diseases caused by fungi, bacteria, and oomycetes lead to substantial losses in agricultural production. Approximately 20% of potential crop yield is lost due to pathogen pressure (Godfray *et al*., 2010). Effective disease management strategies are essential to reduce these losses. In addition to resistance breeding and optimization of crop management practices, application of plant protection products plays a key role in the control of diseases. A major challenge for the application of crop protection products is the development of pathogen resistance, which can progressively reduce their efficacy over time (Hahn, 2014; Massi *et al*., 2021). Furthermore, the application of chemical-synthetic and copper-based compounds has increasingly come under scrutiny, as these substances can have long-term negative effects on the environment and terrestrial and aquatic biodiversity (Pesce *et al*., 2025). In addition, the number of chemical-synthetic active substances and products authorized for crop protection in the EU is decreasing, while the area under organic farming, where the use of chemical-synthetic products is not permitted, continues to increase (Lázaro *et al*., 2021; European Commission, 2025b). Based on these constraints, transnational policy frameworks such as the Farm to Fork Strategy and the European Green Deal call for a substantially reduced application of chemical-synthetic products (European Commission, 2025a). Consequently, there is an urgent need to identify sustainable alternatives that enable the reduction or replacement of chemical-synthetic and copper-based fungicides. In both integrated and organic crop production systems, microbial biocontrol agents (mBCAs) represent an environmentally friendly alternative to chemical control. They consist of bacterial or fungal microorganisms that suppress plant pathogens through either direct activity, such as competition for nutrients or space, antibiosis or parasitism, or through indirect mechanisms, such as the induction of plant defense responses (Köhl *et al*., 2019; Ortiz and Sansinenea, 2021). Due to the limited number of approved mBCAs, their applicability is restricted to a narrow range of crops and pathogens, underscoring the ongoing need for the discovery, development, and regulatory approval of additional mBCAs.

Several species of the bacterial genus *Lysobacter,* including *L. antibioticus*, *L. capsici*, and *L. enzymogenes* have been recognized as biocontrol agents due to their broad antagonistic activity against plant pathogens and nematodes (Ji *et al*., 2008; Tofazzal, 2010; Tang *et al*., 2018; Brescia *et al*., 2021; Martínez-Servat *et al*., 2023; Dinesh Kothari *et al*., 2026). *Lysobacter* species produce a broad range of bioactive compounds, including hydrolytic enzymes, antibiotics, and volatile organic compounds (VOCs), which contribute to their antagonistic effects against plant pathogens (Xie *et al*., 2012; Lin *et al*., 2021). Notably, strains like *L. capsici* AZ78 have shown effective disease control under field conditions, including grapevine downy mildew (Brescia *et al*., 2021), while *L. enzymogenes* C3 has demonstrated broad-spectrum antifungal activity in field trials (Giesler and Yuen, 1998).

We previously characterized *L. enzymogenes* LEC and uncovered its broad-spectrum biocontrol activity (Drenker *et al*., 2023; Drenker *et al*., 2025b). LEC effectively suppressed *Alternaria solani*, *Phytophthora infestans*, *Pseudoperonospora cubensis*, and *Venturia inaequalis* in tomato, potato, cucumber, and apple, respectively. We further identified active metabolites of LEC and characterized their antifungal activity against different pathogens (Drenker *et al*., 2025a).

Building on our findings for diffusible metabolites, we now investigated the activity of volatile compounds (VOCs) from LEC. VOC emission has been identified as a mechanism in biological control, offering potent antimicrobial activity against both bacterial and fungal pathogens while simultaneously enhancing plant systemic resistance (Tilocca *et al*., 2020; Zhao *et al*., 2022). Volatile compounds may complement the activity of metabolites in LEC by broadening the spectrum of activity. The volatile nature of these compounds further enables their diffusion beyond the immediate site of application, potentially reaching plant parts that are difficult to treat with conventional spray applications. Possible applications include closed cultivation systems such as greenhouses, low growing crops with dense canopy and postharvest control.

We conducted a comprehensive characterization of VOC activity of LEC *in vitro* and on cucumber plants, inoculated with downy mildew. This was complemented by characterization of VOC composition using headspace gas chromatography-mass-spectrometry.

Our analyses broaden the understanding of the biocontrol activity of LEC and contribute to the increasing knowledge on the use of microbial VOCs in plant protection.

## 2. Material and Methods

Unless stated otherwise all chemicals and laboratory consumables were obtained from Carl Roth (Karlsruhe, Germany). Media and other solutions were dissolved in deionized water unless specified otherwise. All concentrations given in % refer to (w/v) if not described differently.

### 2.1. Bacterial isolates

The bacterial strains used in this study included *Lysobacter enzymogenes* BI-6432/2 K (hereafter referred to as LEC; Drenker *et al*. (2023)) available in the culture collection of the Julius Kühn Institute, as well as *L. enzymogenes* DSM2043, *L. antibioticus* DSM2044, and *L. gummosus* DSM6980, which were obtained from Leibniz Institute DSMZ (German Collection of Microorganisms and Cell Cultures GmbH). They were cultured alternating on nutrient agar (NA) and soybean flour glucose calcium chloride agar (SGCA; prepared as described in Drenker et al. (2023)) at 25°C for four to five days.

### 2.2. Fungal and oomycetal isolates

The phytopathogenic fungi and oomycetes used in this study were cultured on the media listed in Table 1, as described in Caten and Jinks (1968). Media composition were as follows: rye agar (20%, RA): 20% rye (Alnatura, Germany), 0.5% D(+)-glucose monohydrate, 1.2% agar-agar Kobe; potato dextrose agar (PDA); oatmeal agar (OA, BD DIFCO, Heidelberg), V8 agar: (20% vegetable juice, 0.3% CaCO3, 1.2% agar agar) (Miller, 1955). All fungi and oomycetes were incubated in the dark at 25°C with the exception of *Phytophthora infestans* and *Phoma lingam* (teleomorph *Leptosphaeria maculans*) which were incubated at 15°C. They were transferred onto fresh media every four weeks. The obligate biotrophic oomycete *Pseudoperonospora cubensis* was maintained as a permanent culture on cucumber plants, cultivar Amadora (Beringmeier, Germany).

**Table 1:** Phytopathogenic fungi and oomycotes used in this study. RA: rye agar, PDA: potato dextrose agar, V8: V8 agar, OA: oatmeal agar, JKI: Julius Kühn Institute.

| Fungal Species | Isolate name and origin | Culture medium |
| --- | --- | --- |
| *Alternaria radicina* | A.rad. 1 (JKI, Dossenheim) | 20% RA |
| *Alternaria solani* | A.s. 1 (TU München) | 20% RA |
| *Fusarium graminearum* | 69057 (JKI Berlin) | PDA |
| *Fusarium culmorum* | 283 (Ciba Geigy) | PDA |
| *Phoma lingam* (teleomorph *Leptosphaeria maculans*) | T12aD34 (Universität Göttingen) | PDA |
| *Rhizoctonia solani* | AG-2-2 III B | PDA |
| *Phytophthora infestans* | 606 (Syngenta, Stein, Switzerland) | 20% RA / V8 |
| *Pseudoperonospora cubensis* | not determined (JKI, Dossenheim) | permanent culture on cucumber |
| *Pythium ultimum* | JKI-BI-7202 (JKI, Dossenheim) | OA |

### 2.3. Cultivation of cucumber plants

For the *in planta* trials cucumber plants (*Cucumis sativus*, cultivar Amadora, Beringmeier, Germany) were cultured as described in Drenker *et al*. (2023) with the following modifications: The substrate was put into a drying cabinet at 60°C for at least two days to eliminate fungus gnats’ eggs. Afterwards it was stored in a closed plastic bin and moisturized before further usage. After transplanting the seedlings into individual pots, the pots were placed in trays and irrigated from below with 1 L of water per tray every Monday, 0.5 L every Thursday, and 1 L of a 0.2% (v/v) Hakaphos Blau fertilizer solution (Compo Expert, Germany) every Friday. Plants were grown under fluorescent lights (Philips Master TL-D 36W/840, Netherlands) with roughly 80 cm distance to the plants (PAR: ca. 100 µmol/m2s; light regime 16:8 h, 25°C).

### 2.4. Influence of VOCs on pathogen growth

To identify a medium for optimal VOC production by LEC the media NA, TSA and SGCB were tested. The inhibitory effect on the growth of plant pathogens *A. radicina* and *R. solani* was analyzed in split-plate assays carried out as follows: Petri dishes with a barrier in the middle were filled with 12.5 mL of the respective medium to test on one side and 12.5 mL of rye agar (RA, 10%) on the other. In parallel a 1:10 dilution of the respective LEC growth medium was tested to simulate starvation conditions. Subsequently, 50 µL of 10×7 cells/mL of LEC in 0.6% NaCl were spotted on the LEC growth medium compartment of the split plate using a flame sterilized loop while keeping a 5 mm distance to the edge and central barrier to prevent LEC to reach the other side of the Petri dish. Petri dishes were sealed with parafilm and incubated for three days at 25°C in the dark. Subsequently, an agar plug (7 mm diameter) was cut from the border of a seven-day old fungal colony using a sterile cork borer and placed on the RA compartment of the split plate next to the center of the barrier. Petri dishes were sealed with two layers of parafilm and incubated at 25°C in the dark with the lid showing upwards. Petri dishes containing a fungal plug exclusively served as control. Mycelial growth was assessed by measuring the growth radius from the plug center at 45°, 90° and 135° relative to the plastic barrier when the corresponding control reached the edge of the Petri dish.

To analyze the inhibitory effect of VOCs from LEC grown on NA medium on the growth of additional plant pathogens, split-plate assays were carried out as described before. For this assay mycelial growth was assessed daily. Three to five Petri dishes were analyzed for each treatment as indicated in the respective figure legend (Fig. 1) and the experiment was repeated at least three times per pathogen.

**Figure 1.**
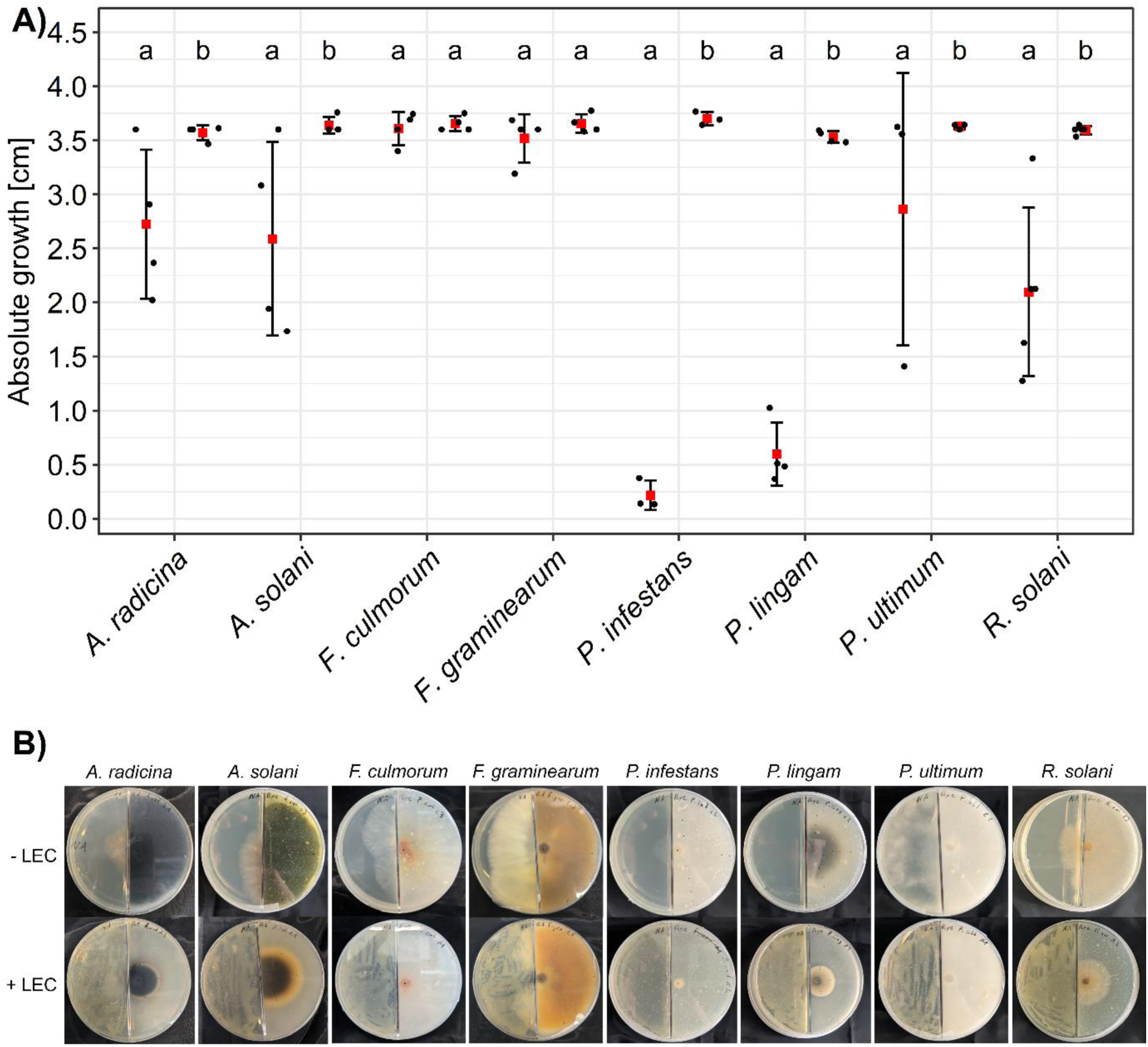
Influence of volatile organic compounds (VOCs) of *Lysobacter enzymogenes* LEC on growth of six fungi and two oomycetes. A) Growth reduction was determined in split-plate assays and was measured in three positions per Petri dish with three to five Petri dishes per treatment. Absolute growth in cm for each treatment (left) and respective control (right) is shown; *A. radicina*: *Alternaria radicina* (n = 4), *A. solani*: *Alternaria solani* (n = 4), *F. culmorum*: *Fusarium culmorum* (n = 4), *F. graminearum*: *Fusarium graminearum* (n = 4), *P. infestans*: *Phytophthora infestans* (n = 3), *P. lingam*: *Phoma lingam* (n = 4), P. ultimum: Pythium ultimum (n = 3), R. solani: *Rhizoctonia solani* (n = 5). Red box: mean, jitter-dots: individual experimental repetitions, error-bars: standard deviation. p-value < 0.05, data analysis: Generalized linear mixed model, post-hoc analysis: estimated marginal means (EMMs) with pairwise comparisons between control and treatment, p-value adjustment: Tukey method. B) Representative presentation of a split-plate assay for all tested fungi and oomycetes with (+ LEC) and without (-LEC) *Lysobacter enzymogenes* LEC VOC treatment on the left side of the split-plate and the fungal/oomycetal isolates on the right side. Shown are representative Petri dishes after seven days (*A. solani*), eight days (*A. radicina*), four days (*F. culmorum*), five days (*F. graminearum*), nine days (*P. infestans*), ten days (*P. lingam*), three days (*P. ultimum*), four days (*F. culmorum*), and three days (*R. solani*) incubation at 25°C. A. solani, A.radicina, F. culmorum, F. graminearum, *P. infestans*, *P. ultimum* and *R. solani* were partially overgrown to the left side on the control plates (-LEC).

For the analysis of additional ventilation on the inhibitory effect of LEC VOCs the experiments were carried out as described above with the following changes: After the fungal plugs were added and the plates sealed with parafilm, four slits (1 cm wide) were cut into the parafilm, two on the LEC and two on the pathogen side at an 45° angle left and right from the barrier. As a control plates without slits were used for all treatments as well as plates with and without LEC respectively as described above as a control. The experiment was done two times with three to four plates for each repetition. At the end of both experiments the OD600 was determined as an approximation for cell density to assess the influence of a possibly varying cell number due to changed culturing conditions compared to the non-ventilation conditions using a standard photometer. The respective suspension was prepared by scraping the whole amount of LEC cells from a single plate by pipetting 3 mL of a 0.6% NaCl solution onto the plate and using a spatula to scrape them of. This was repeated two more times. The suspension was collected in a 15 mL falcon and filled up to 10 mL. The obtained suspensions were diluted 1:10 in a 0.6% NaCl solution for the measurement. Relative growth inhibition was calculated as follows:

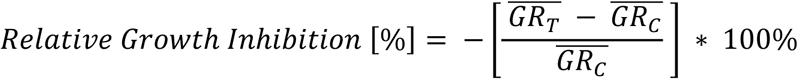

where: *GR^-^_c_*: mean growth radius of untreated pathogen (without exposure to LEC VOCs). *GR^-^_T_*: mean growth radius of treated pathogen. The radius of the plug (0.3 cm) was subtracted from each measurement.

### 2.5. Influence of VOCs on pH

To determine the influence of the VOCs on the pH of the agar, LEC was spread exclusively on split Petri dishes as described above. After 72 hours the pH on the opposing side (10% RA) was measured by placing a strip of pH indicator paper (pH-Fix pH 4.5 - 10) on the agar. As a control, plates without LEC were used. Three replicates (Petri dishes) were used, the experiment was repeated three times.

### 2.6. Analysis of VOC phytotoxicity on cucumber plants

To determine if LEC VOCs show any phototoxic effects on cucumber plants, cultivar Amadora were inoculated with *P. cubensis* as described in (Drenker *et al*., 2023) with the following modifications: After inoculation each plant was kept in a separate plastic propagator (Meyer Rellingen, Germany) and immediately a square Petri dish (12×12 cm) with LEC on NA was placed without lid below the plant into the plastic propagator to mediate VOC distribution into the plastic box. Each Petri dish was previously spot inoculated with 100 µL of a LEC suspension (10×7 cells/mL), sealed with parafilm and incubated at 25°C for 72 h in the dark. To ensure high relative humidity and to mimic inoculation conditions (described in section 2.7.) the leaves were sprayed with well water on the adaxial side, the lids of the plastic propagators were sprayed on the inside, and all ventilation openings were closed and sealed with tape. The plants were transferred into a greenhouse chamber and incubated for five days with the propagator lids closed. The plants were watered every three days with 30 mL well water. Two plants per treatment were analysed for phytotoxicity by monitoring all visible changes in plant appearance. As a control treatment a non-inoculated Petri dish was placed below the plants as described above. Effects were assessed one, two, three, and five days after inoculation.

### 2.7. Activity of LEC VOCs against *P. cubensis* on cucumber plants

The experiment was carried out as described in Section 2.6 with the following adjustments: Immediately after inoculation of the plants with *P*. *cubensis* a Petri dish with 60 mm diameter with LEC on NA was put below the plants into the plastic propagator. Each Petri dish was previously spot inoculated with 20 µL of a LEC suspension (10×7 cells/mL), sealed with parafilm and incubated at 25°C for 72 h. The plants were then put into three individual GroBank compartments (CLF PlantClimatics, Germany) according to their treatment to prevent any cross-over effects of VOCs. After 24 h the ventilation slits were opened and precipitated water was removed from the propagator lids with a tissue paper. Subsequently, the plastic propagator lids were put back on. The plants were then incubated at a 16:8 hour light:dark regime under the following conditions: light 100 µmol m^-2^ s^-1^ and 21°C, 60% relative humidity and 16°C and 75% relative humidity at night for seven days. The lids were left on to assure exposure to the VOCs. The plants were watered every three days with 30 mL well water using a tube and a syringe to keep the lids on during the process. Ten plants per treatment were used, and the experiment was repeated twice. Four plants were sprayed with water solely and used as a healthy control in each experiment. As no adverse effects were observed during the previous screening from plates containing NA only, no such control was included in subsequent experiments. The relative symptomatic leaf area was assessed as described (Drenker *et al*., 2023).

### 2.8. Identification of a non-inhibitory strain as GC-MS negative control

To gain insight into which VOC compounds contribute to the disease-suppressive effect of LEC, a *Lysobacter* isolate without disease-suppressive activity was screened for. To do so, *L. enzymogenes* DSM2043, *L. antibioticus* DSM2044 and *L. gummosus* DSM6980 were tested regarding their inhibitory potential against the pathogens with the strongest sensitivity in the previous tests *R. solani*, *P. infestans* and *P. lingam* in split Petri dish assays. The assays were carried out as described in section 2.4 for LEC. The respective *Lysobacter*-pathogen combinations were prepared in triplicates and the experiment was done once.

### 2.9. VOC sampling and analysis by headspace analysis

Headspace from both *L. enzymogenes* cultures and growth media was carried out using a Turbomatrix-40 Trap (PerkinElmer, Germany). Bacterial cultures were directly grown in 22 mL headspace vials (PerkinElmer, Germany) filled with 8 mL nutrient agar. During solidification of the agar the vials were inclined 12° upwards to maximize the agar’s surface area. For the inoculation 50 µL of a 10×7 cells/mL suspension was used.

The vials were left without a cap under the sterile bench until the excess liquid evaporated. Subsequently the vials were kept in the inclined position with the cap on the opening without sealing them and incubated at 25°C for 48 h in the dark. The vials were closed airtight after 48 h of bacterial growth, 24 h before the headspace analysis. The vials were thermally regulated at 70°C in the headspace oven with shaking for 5 min. Thereafter, the vials were pressurized for one minute with 240 kPa of helium followed by pressure release for 1.2 minutes collecting the vapor phase from the vial via the sample syringe on the absorbent trap. The needle and the transfer line were heated to 75°C and 80°C, respectively. Sampling was repeated three times per vial. Adsorbed water was removed by dry purging the trap for five minutes. The focused analytes on the trap were released spitless into the GC system by heating up the trap to 280°C. Trap desorb pressure was set to 60 kPa and the column pressure was regulated at 150 kPa through the analysis by the headspace sampler. The headspace trap system was directly on-column connected through an unpacked capillary transfer line to a GC-2400 coupled to a MS-2400SQT (PerkinElmer, Germany). The GC was equipped with a nonpolar Elite-5MS capillary column (30 m × 0.25 mm id × 0.25 µm film thickness, PerkinElmer, Germany). The oven temperature was programmed as follows: 40°C for 1 min, heating from 50 to 180°C at 5 K/min, increased to the final temperature of 280°C by a rate of 20 K/min, and held at 280°C for six minutes. The GC inlet line temperature and the ion source temperature were set at 250°C and 180°C, respectively. The quadrupole mass detector was operated in the electron impact (EI) mode at 70 eV. All data were obtained by collecting the full-scan mass spectra within the range of 50–550 m/z. Three samples of each bacterial strain and two growth media samples were analyzed within a day for three days, resulting in eight - nine samples per bacterial strain and six growth media samples. For quality control and the determination of retention indices (RIs), an alkane standard mixture (C8–C20) was analyzed using the same GC-MS method. Additionally, the NIST-based identification of *p*-xylene, benzaldehyde, decane, phenylacetaldehyde, acetophenone, and *n*-dodecane was verified using authentic reference standards.

Volatile compounds from the chromatograms were identified and quantified with the “Automated Mass spectral Deconvolution and Identification System” (AMDIS, V. 2.71; National Institute of Standards and Technology NIST, Boulder, CO) following previously described methods (Gross *et al*., 2019), in combination with the NIST MS Search program (Version 2.0) and the NIST/EPA/NIH Mass Spectral Library (NIST 08). Retention indices were calculated based on the n-alkane series (C8–C20) and used for retention index data analysis with AMDIS. The deconvolution was applied with the following settings: component width, 32; adjacent peak subtraction, one; resolution, low; sensitivity, low; shape requirements, medium. For Identification match factor was set to 70 and relative retention index deviation ≤ 5% + 0.02 from the reference value and application of average match factor penalties: Max. penalty: 20 and “no RI in library”: 10. Components with a signal to noise ratio < 20 were excluded from the analysis. Tentatively identified compounds were assigned by comparison with the NIST mass spectral library. Compounds that could not be identified were labeled according to their respective RI values to allow comparison between samples. Pyrazine-like compounds were identified based on their characteristic mass spectral patterns; however, the positions of substituents (positional isomers) were not determined.

### 2.10. Analysis of cell morphology by microscopy

To investigate the effects of VOCs on *R. solani* in split Petri dishes as described in section 2.4 with three replicates per treatment. Analyses were done once the *R. solani* control treatment without exposure to LEC VOCs reached the edge, in comparison to VOC-treated tissue of *R. solani*. The youngest mycelial tissue, from the edge of the colony was analysed. Number of repetitions and analysed samples are described below for fluorescence and transmission electron microscopy (TEM).

#### 2.10.1. Fluorescence microscopy of *R. solani*

Double staining with DAPI and Nile Red was performed on mycelial tissue of *R. solani* to visualize nuclei (DAPI) and intracellular lipid structures (Nile Red). DAPI stock solution (0.1 mg/mL) was prepared in ultrapure water and stored in the fridge. The working solution (0.1 µg/mL) was obtained through a dilution in phosphate-buffered saline (PBS), pH 7.4. Nile Red stock solution (500 µg/mL) was prepared in acetone (99.5%) and stored at −20°C. The working solution (5 µg/mL) was prepared by dilution in PBS, pH 7.4. Samples were prepared on adhesive microscope slides (Superfrost Plus Gold; Thermo Scientific, Germany) using PBS, pH 7.4. For sample preparation, mycelial tissue was carefully transferred with a toothpick by gently sliding over the top of the fungal colony and transferring it into a drop of PBS on a microscope slide, followed by gentle dispersing to remove excess liquid. After brief air-drying, samples were incubated with the DAPI working solution for 5 min at room temperature in the dark, followed by two washes with PBS, pH 7.4. To do so, DAPI was removed by holding a piece of filter paper (absorption paper MN224, Macherey-Nagel, Germany) against the border of the microscope slide. Subsequently, samples were incubated with the Nile Red working solution for five minutes and washed twice with PBS as described above. Coverslips were placed on the samples, which were then incubated for 24 hours in a humid chamber to ensure homogeneous penetration of both dyes throughout the mycelial tissue. Fluorescence microscopy was performed sequentially for each fluorophore with the Zeiss Axio Imager M2 microscope using the ZEN blue software (version 3.3). The settings were as follows: DAPI was visualized by excitation at 385 nm and fluorescence emission was detected with the 96 HE BFP filter. Nile Red was excited at 555 nm, and emission was detected using the 43 HE Cy3 filter. Samples were analysed from three different border regions per plate and three plates per treatment. The experiment was repeated three times.

#### 2.10.2. Transmission electron microscopy (TEM) of *R. solani*

Mycelial tissue was fixed in Karnovsky’s fixative (2.5% glutaraldehyde and 2% paraformaldehyde in 0.1 M cacodylate buffer, pH 7.4) at 4°C for two hours. After fixation, samples were washed three times for 30 minutes in 0.1 M sodium cacodylate buffer (Na(CH₃)₂AsO₂·3H₂O), followed by incubation in 2% osmium tetroxide (OsO₄) in 0.1 M sodium cacodylate buffer for two hours. For contrast enhancement, samples were stained with 2% uranyl acetate in 70% ethanol for two hours. Dehydration was performed using a graded ethanol series consisting of 70% (1 h), 80% (1 h), 96% (1 h), and 100% ethanol (3 x 1 h). Embedding for TEM analysis was conducted using propylene oxide and Spurr resin (Sigma-Aldrich, USA). Dehydrated samples were first incubated in propylene oxide (2 x 30 min), followed by gradual infiltration with increasing concentrations of Spurr resin: propylene oxide/Spurr ratio 3:1 (1 h), 1:2 (1 h), 1:3 (1 h), and pure Spurr resin (2 x 3 h). Polymerization was achieved by heating the samples at 70°C for eight hours. Ultrathin sections (50-70 nm) were cut using an ultramicrotome (PowerTome XL, RMC Products by Boeckeler Instruments, Inc., USA). Sections were contrasted with uranyl acetate for ten minutes and rinsed with double-distilled water. Subsequently, lead citrate staining was performed for two minutes; NaOH pellets were used to absorb atmospheric CO₂ and prevent precipitate formation. After a final wash with double-distilled water, sections were examined in a Zeiss TEM 900 microscope (Zeiss, Germany) and imaged with a CCD camera system using the Image SP software (Sysprog & TrS, Belarus).

### 2.11. Statistical analysis

The data sets were statistically analyzed using a generalized linear mixed model (glmm) approach. To do so, R (4.4.0) (R Core Team, 2024) and RStudio (RStudio 2023.03.0+386 “Cherry Blossom”) were used together with the R packages glmmTMB (1.1.12) (Brooks *et al*., 2017) for model building, DHARMa (0.4.7) (Hartig, 2024) for model diagnostics, emmeans (2.0.1) (Lenth, 2025) for post-hoc analysis (α = 0.05) applying Tukey’s correction for multiple comparisons and ggeffects (2.3.2) (Lüdecke, 2018) to calculate the 95% confidence interval (95% CI).

In brief, the respective response variable (mean “Absolute growth [cm]” per pathogen and mean “Relative infected leaf area [%]” per plant or per leave, respectively) was fitted against the respective treatment including the experimental repetition as a random-effect (+ (1 | run)) to account for variance due to the experimental repetition. The volatile data was fitted using a negatively skewed normal distribution based on the data histogram showing a tendency for a ceiling effect due to the growth limit set by the size of the Petri dish (maximum growth). For the *in vitro* data the treatment (Ctrl) and for the *in planta* data the experimental repetition (Run) was included as a dispersion parameter to account for additional variance between treated and untreated pathogens and between the experimental repetition respectively. For the *in vitro* data an interaction term was included (Ctrl * Fungi) to account for the different degrees of susceptibility of the different pathogens to the volatiles. For the *in planta* data a nested random effect was included to account for plant specific variance between experimental repetitions and to account for non-independence of plants and hence for pseudo-replication (+ (1 | Run:Plant). A ziformula was included to account for excess zeros (real biological zeros) from uninfected control plants. Additionally, the *in planta* data was transformed from the percent scale (1 – 100%) to an interval [0,1] in order to fit an ordbeta model based on the data histogram showing data points accumulating at values 0 and 1. The models were analyzed regarding their quality of fitting the experimental data. Therefore, the distribution of the residuals, potential outliers, and violations of the normality and homoscedasticity assumption for the residuals was checked using the DHARMa library by plotting the simulated output. Potential over-and under-dispersion of the fitted models was tested using testDispersion(). Potential outliers were analyzed using testOutliers() and outliers(). The 95% confidence interval was calculated using the function predict_response() (Fig. S1, S2, Table S1, S2).

Differences in VOC composition were assessed using PERMANOVA with 10,000 permutations based on robust Aitchison distances calculated from relative abundances (% of total peak area), as implemented in the vegan package (Oksanen *et al*., 2024). This approach accounts for the compositional nature of the data and the presence of zero values. Pairwise comparisons were conducted using the pairwise.perm.manova function from the RVAideMemoire package (Hervé, 2023), with p-values adjusted for multiple testing using the Bonferroni correction. Homogeneity of multivariate dispersions among groups was assessed using the betadisper function in the vegan package, followed by analysis of variance (PERMDISP). Non-metric multidimensional scaling (NMDS) based on robust Aitchison distances was used to visualize differences in VOC compositions. To assess differences in the emission of individual VOCs, absolute peak areas were compared among treatments using Kruskal-Wallis tests. Absolute peak areas were used to reflect compound abundance and to compare emission levels between treatments. For VOCs showing significant differences in Kruskal-Wallis tests (FDR-corrected), pairwise comparisons were performed using Dunn’s test with Benjamini-Hochberg correction. For visualization a heatmap was generated from mean absolute peak areas per treatment, and values were log₁₀(x + 0.01)-transformed to account for zero values and improve visualization. The raw data and corresponding R scripts are available in a GitHub repository (https://github.com/JulianMaierJKI/L.enzymogenes_Volatile_data).

## 3. Results

### 3.1. LEC VOCs inhibit the growth of pathogenic fungi and oomycetes

In order to identify a suitable growth medium for subsequent VOC analyses, LEC was grown on different media and at different media concentrations. Subsequently the suppressive activity of the released VOCs was determined in split-plate assays by assessing the inhibition of mycelial growth of *A. solani* and *R. solani*. VOC activity was strongest when LEC was grown on NA or TSA medium, (Fig. S3, Table S3), in comparison to lower concentrated 10% NA, 10% TSA and to SGCA medium of both concentrations. All subsequent experiments were therefore done with LEC propagated on NA medium, which further facilitated comparability to previously published studies for *Lysobacter* spp. (Lazazzara *et al*., 2017).

Fungi and oomycetes showed varying degrees of growth inhibition upon exposure to LEC VOCs (Fig. 1A, B). Mycelial growth of *P. infestans and P. lingam* was significantly inhibited when exposed to LEC VOCs, with a growth radius of only 0.22 cm and 0.6 cm compared to around 3.5 cm for the control treatments, respectively (Fig. 1A). The growth radius for *R. solani* was 2.1 cm,2.59 cm for *A. solani* and 2.72 cm for *A. radicina.* Therefore, their inhibition was less but still significantly when compared to their respective control. It has to be noted, that although *P. ultimum* growth was significantly different compared to the control treatment it was only inhibited in one trial out of three repetitions (1.4 cm). Contrary to that, growth of *F. culmorum* and *F. graminearum* was not significantly affected by LEC VOCs. The observed growth effects did neither depend on pathogen growth dynamics (e.g. fast growers *P. ultimum* and *R. solani* and slow growers *A. radicina* and *P. lingam*) (Fig. 1A) nor their taxonomic affiliation, e.g. fungi and oomycetes. The observed growth showed a gradual and consistent progression over time without abrupt changes (Fig. S4).

Since *Lysobacter* spp. are described as aerobic organisms (Reichenbach, 2006) we investigated the influence of ventilation by adding slits into the parafilm covering the Petri dishes. It led to drastic increases in growth inhibition for *A. radicina*, *A. solani* and *R. solani* by over 95%, compared to suppression under sealed conditions, 4.1%, 3.5% and 35% respectively (Fig. S5, Table 2). In addition, the resulting optical density (OD600) as a proxy for cell density indicated that the observed increase in inhibition was not due to an increase in cell number. The pH value of RA was determined with and without VOC exposure to assess whether fungal growth alterations are associated with changes in pH. An average increase of pH (7.3 ± 0.25) as compared to the non-treated control (6.2 ± 0.29) was observed (Table S4).

**Table 2:** Effect of increased ventilation on radial growth inhibition of fungal pathogens by VOCs of *Lysobacter enzymogenes* LEC and on LEC biomass concentration. Radial fungal growth (%) was determined in split-plate Petri dishes in three positions per Petri dish with three to four Petri dishes per treatment. Shown are mean growth reduction values determined in relation to a non-treated control with and without added ventilation slits in the surrounding parafilm. *A. radicina*: *Alternaria radicina*, *A. solani*: *Alternaria solani*, *R. solani*: *Rhizoctonia solani*. LEC biomass was determined simultaneously by OD600 measurement for two independent experimental repetitions except for *A. radicina* with one experimental repetition.

| Fungi | Ventilation | Growth reduction [%] | Mean OD600 |
| --- | --- | --- | --- |
| *A. radicina* | - | 4.1 | 0.214 |
| *A. radicina* | + | 95.8 | 0.048 |
| *A. solani* | - | 3.5 | 0.281 |
| *A. solani* | + | 95.5 | 0.050 |
| *R. solani* | - | 35.0 | 0.351 |
| *R. solani* | + | 95.8 | 0.131 |

### 3.2. LEC volatiles decrease pathogen infection *in planta*

Transferability of VOC-induced inhibition *in vitro* to crop plants was investigated using cucumber plants inoculated with downy mildew (*Pseudoperonospora cubensis*) due to their short growth height and the broadness of their leaves. To avoid phytotoxic effects (Fig. S6), 20 µL of LEC suspension were used for all experiments. Leaves of infected control plants showed a relative infected leaf area of 95% (Fig. 3, CON+). Leaves situated directly above Petri dishes with LEC cultures and directly exposed to VOCs had a mean infection of 39% (Fig. 2, +LEC volatiles). Leaves adjacent to the Petri dish with LEC (-LEC volatiles) had disease levels of about 90%, but still significantly lower than the diseased control. However, both treatments were not as effective as the treatment with the chemical standard Cuprozin Progress (0.52%).

**Fig. 2.**
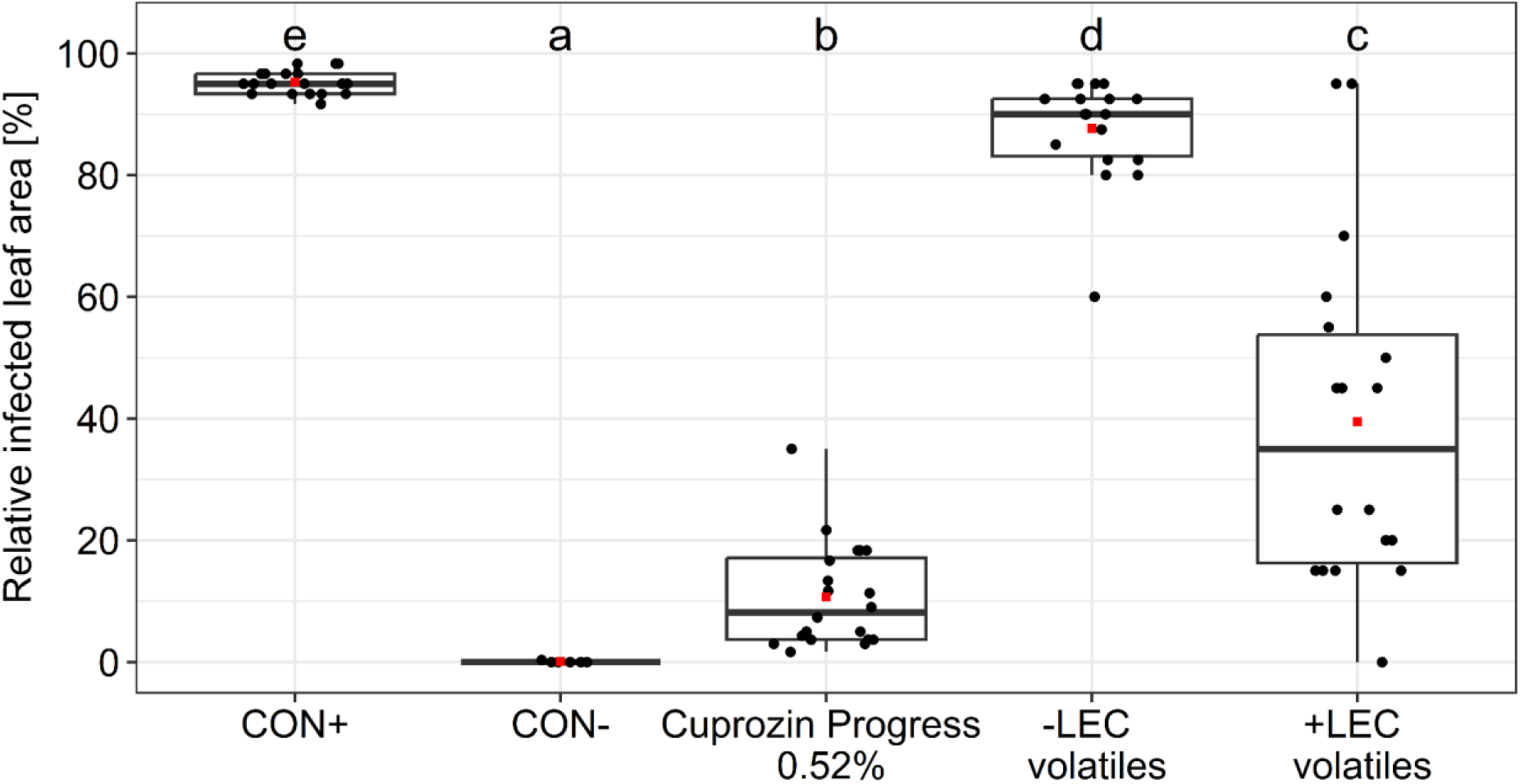
Effect of VOCs of *Lysobacter enzymogenes* LEC on disease severity (relative infected leaf area in %) of *Pseudoperonospora cubensis* on cucumber cultivar Amadora. CON+: diseased plants, inoculated with *P. cubensis* (n = 20 plants, three leaves per plant), CON-: healthy plants (n = 7, three leaves per plant), Cuprozin progress 0.52%: plants treated with chemical fungicide (n = 20, three leaves per plant), -LEC volatiles: leaves adjacent to the Petri dish with LEC (n = 18, two leaves per plant), +LEC volatiles: leaves immediately above the Petri dish with LEC (n = 18, one leaf per plant), n: plants per treatment. Due to a strong dependency on proximity of the VOC-source to the infected leaf, leaves directly above Petri dishes were analyzed separately from VOC-source adjacent leaves. Each data point represents the mean relative infected leaf area (%) based on the indicated number of leaves per plant. p-value < 0.05 based on generalized linear mixed model, post-hoc analysis: estimated marginal means (EMMs), p-value adjustment: Tukey method. Different letters indicate statistically significant differences. Horizontal box-plot line: median, red box: mean and jitter-dots: individual values per plant (leaf(s)) and per treatment.

**Fig. 3.**
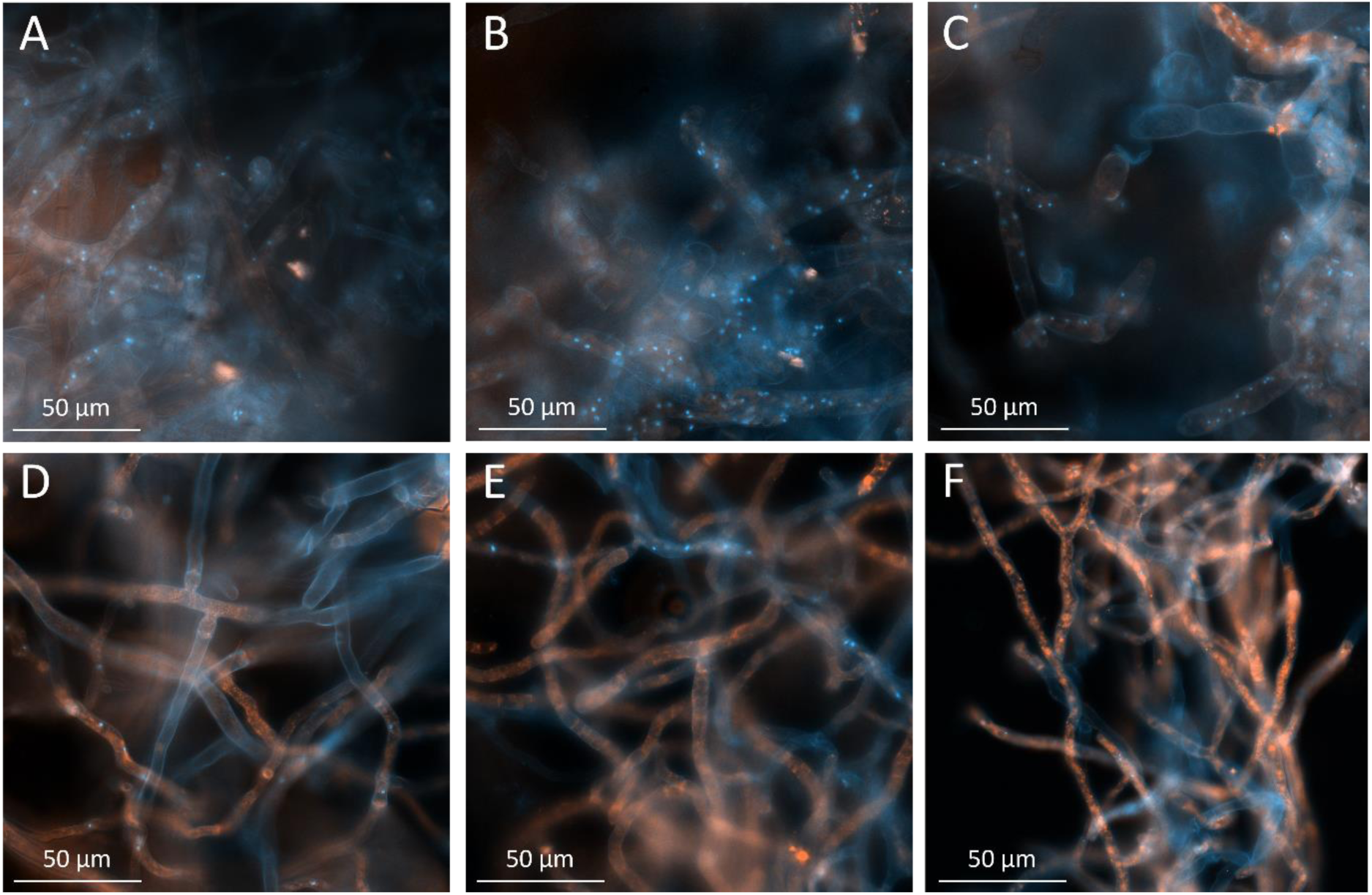
Effects of volatile organic compounds (VOCs) of *Lysobacter enzymogenes* LEC on nuclei and lipid abundance of *Rhizoctonia solani*, analysed by fluorescence microscopy. The fungus was grown for three days in split Petri dishes either in the absence (A-C) or presence of LEC VOCs (D-F). Subsequently, mycelial tissue from the actively growing hyphal margin was stained with DAPI and Nile Red. DAPI stained nuclei as blue dots, whereas lipids are stained in red by Nile Red. Images were acquired with the Zeiss Axio Imager M2 microscope and the ZEN blue software (version 3.3). Ten sections were analysed per treatment, the experiment was repeated three times.

### 3.3. Cellular morphology is altered upon VOC treatment

Mycelial tissue of *R. solani* was stained with DAPI to visualize nuclei and Nile Red to detect lipids. Both were chosen in order to detect cell stress on a cellular level. Considerable variability in cell morphology, nuclear abundance and lipid distribution was observed between samples and among neighbouring cells. Consistent differences between the nontreated control and the VOC treatment became apparent when evaluating ten samples per treatment in three replicates.

In non-treated mycelia multinucleate cells were observed more frequently (blue dots in Fig. 3A-C), whereas VOC-treated mycelia (Fig. 3D-F), showed a higher occurrence of cells lacking visible nuclei and fewer multinucleate cells overall. Contrary, lipid accumulation seemed to be enhanced in the VOC-treated cells, as indicated by the intense and abundant Nile Red staining (Fig. 3D-F).

In addition, transmission electron microscopy revealed differences in lipid appearance and morphology between treatments. In nontreated cells, lipid bodies displayed sharp boundaries and a well-defined appearance (Fig. 4A, C), while lipid bodies in VOC-treated cells appeared more diffuse and less clearly delineated (Fig. 4B, D). Taken together, VOCs of LEC affected nuclei and lipid abundance of *R. solani,* while other cell structures were not obviously affected.

**Fig. 4.**
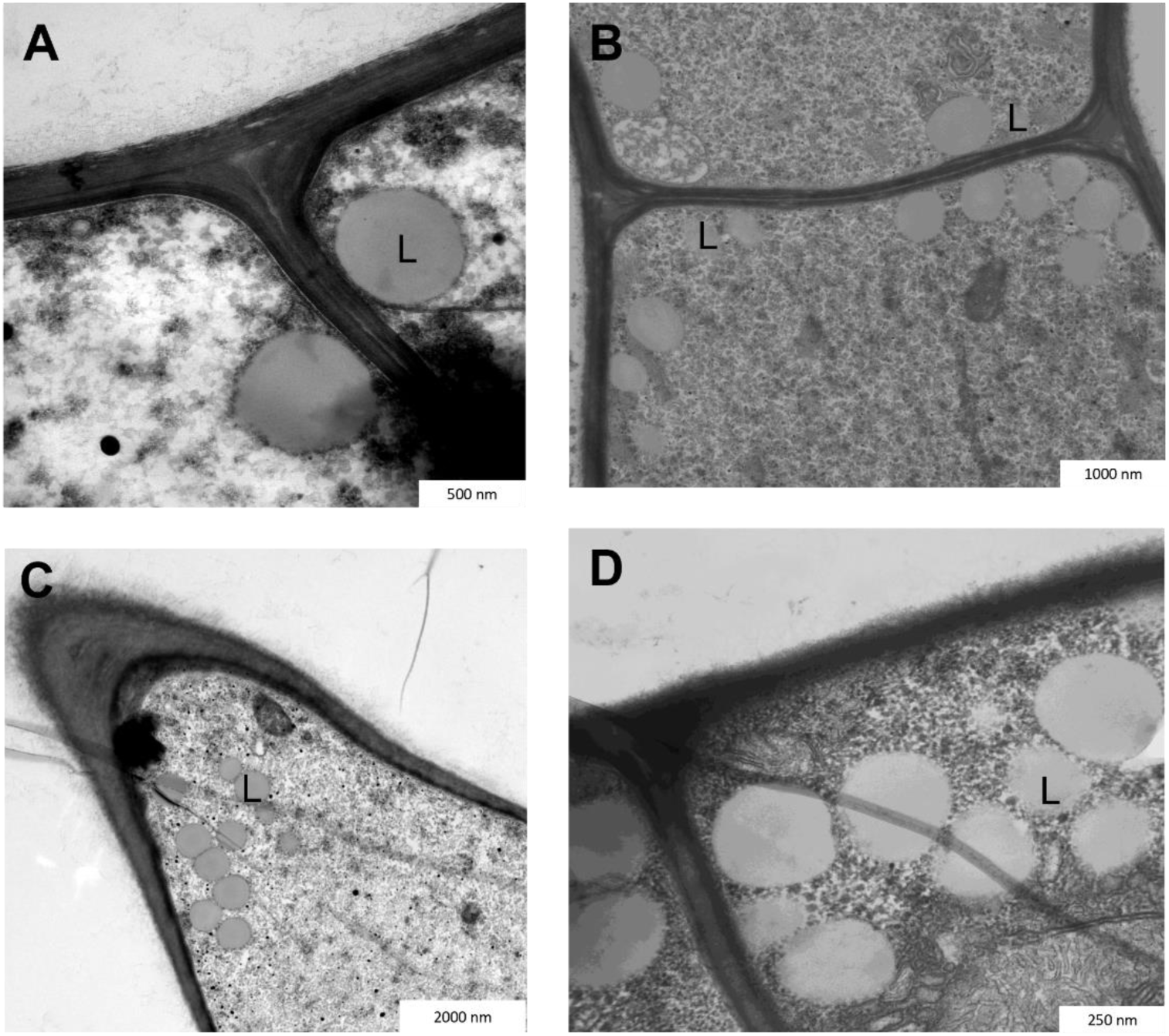
Ultrastructural analysis of Rhizoctonia solani mycelial tissue upon exposure to volatile organic compounds (VOCs) of Lysobacter enzymogenes LEC, examined by transmission electron microscopy. The isolate was grown for three days in split Petri dishes, with (B, D) or without (A, C) exposure to LEC VOCs. Subsequently actively growing hyphae were fixed, contrasted, and sectioned. Images were acquired using a transmission electron microscope and Image SP software. (L) indicates representative lipid bodies within R. solani cells.

### 3.4. VOC profile differs between closely related *L. enzymogenes* strains

To identify the VOCs which are potentially responsible for the observed growth suppression, the VOC profiles of LEC and *L. enzymogenes* DSM2043 were compared. The latter one was chosen based on previous analyses, where we searched for a strain, that was taxonomically closely related to LEC, but at the same time lacking inhibitory effects against *R. solani* (Fig. 5, Table S5). Volatile substances detected in LEC, but not in DSM2043 might be involved in its biocontrol activity.

**Fig. 5.**
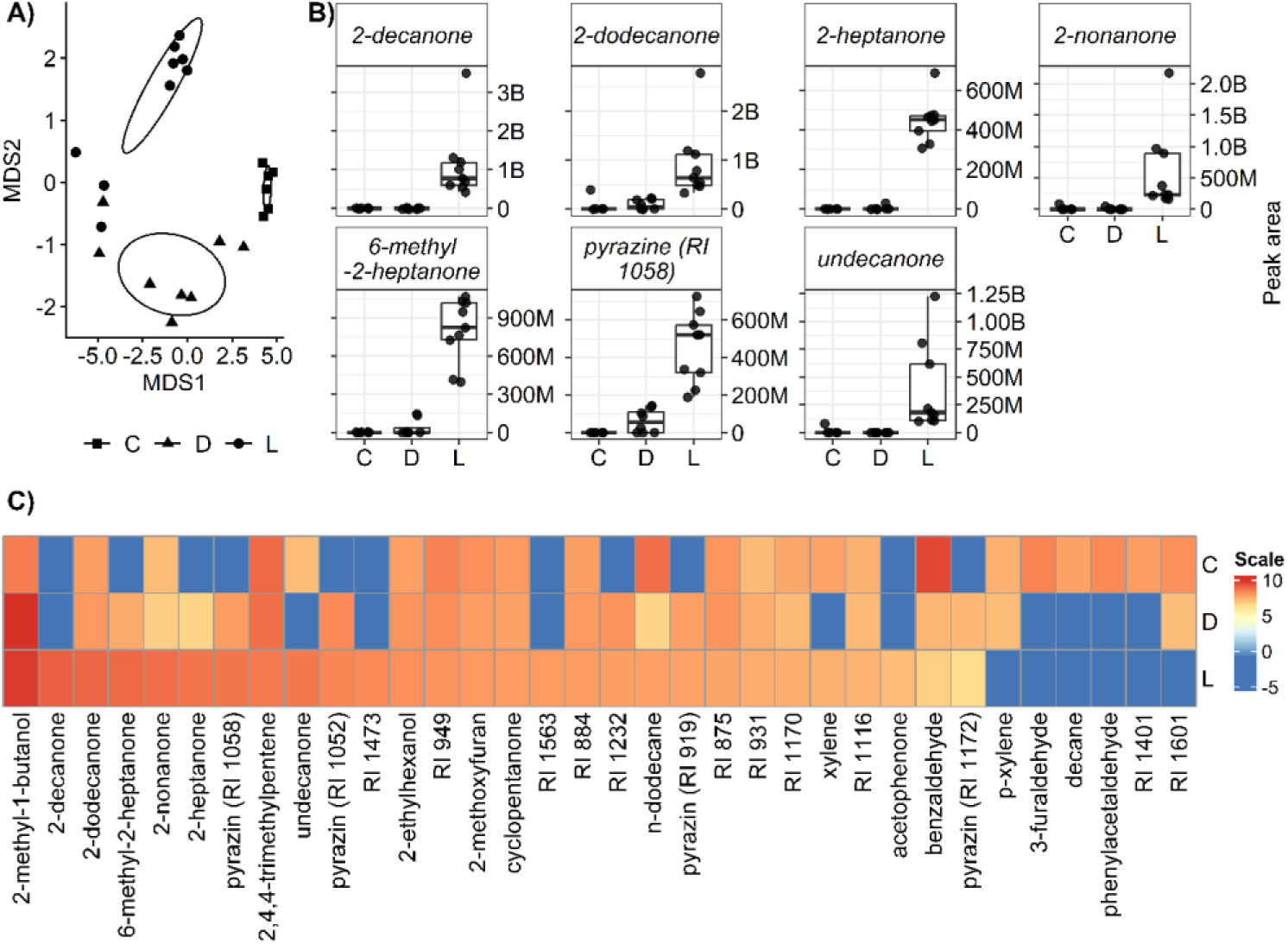
Volatilome analysis of *Lysobacter enzymogenes* LEC, *L. enzymogenes* DSM 2043, and pure growth medium. A) Non-metric multidimensional scaling (NMDS) plot based on robust Aitchison distances calculated from proportional volatile compositions (2d-stress: 0.127). B) Emission of selected volatile organic compounds (VOCs) specifically abundant in *L. enzymogenes* LEC compared to DSM2043 and the growth medium. Boxplots represent the distribution of absolute peak areas across replicates, with individual data points overlaid. VOCs were selected based on significant differences between LEC and both DSM2043 and the medium (Dunn’s test, Benjamini–Hochberg adjusted p ≤ 0.05), C) Heatmap of emitted VOCs. Values represent log₁₀-transformed mean peak areas across replicates. VOCs were clustered based on Euclidean distance to reveal similarity patterns, while treatments were kept in a fixed order. The heatmap highlights compounds higher abundant (red) or lower abundant (blue) in LEC (L: n=9) compared to DSM2043 (D: n=8) and the control medium (C: n=6).

In total, 34 volatile components (Fig. 5C) were annotated from the chromatograms and the overall volatile composition differed significantly between both strains and the control medium (Fig. 5A, PERMANOVA, F = 8.026, R² = 0.445, p <0.001, df = 2). No significant differences in multivariate dispersion were detected among groups (PERMDISP/ANOVA, p = 0.111), indicating that PERMANOVA results are not driven by differences in within-group variability. Pairwise PERMANOVA revealed significant differences between the control medium and DSM2043 (p = 0.0015), the control medium and LEC (p = 0.0018), as well as between the two bacterial strains (p = 0.0036; Bonferroni-adjusted). (Fig. 5A). LEC showed significantly higher peak areas for 2-decanone, 2-dodecanone, 2-heptanone, 2-nonanone, 6-meythy-2-heptane, pyrazine and undecanone as compared to DSM2043 and the control medium (Fig. 5B, C, Table S6). In addition to these differences in ketone abundances, several pyrazine-like compounds were detected in the VOC profiles of both strains but not in samples from the growth media (Fig. 5C). A compound with RI 919 showed a molecular ion at *m/z* 108 and characteristic fragment ions at *m/z* 95, 81, 107, and 109, consistent with a dimethylpyrazine. Compounds with RI 1052, 1058, and 1172 exhibited molecular ions at *m/z* 138 and characteristic fragment ions at *m/z* 137, 120, 109, and 107, consistent with tetramethyl-substituted pyrazines.

## 4. Discussion

Microbial biocontrol agents are a promising alternative to chemical-synthetic fungicides for sustainable crop production. In this context we previously identified and characterized *L. enzymogenes* LEC as a biocontrol agent with suppressive activity against several agronomically important fungi and oomycetes and determined diffusible metabolites related to biocontrol activity in the culture broth (Drenker *et al*., 2023; Drenker *et al*., 2025a; Drenker *et al*., 2025b). We now characterized volatile organic compounds (VOCs) and the antimicrobial activity of LEC.

We demonstrate that LEC produces VOCs active against a range of phytopathogenic fungi and oomycetes. This is in agreement with previous studies (Garbeva *et al*., 2014; Asari *et al*., 2016; Lazazzara *et al*., 2017; Vlassi *et al*., 2020; Rani *et al*., 2023), where inhibitory effects were observed for *P. infestans* and *R. solani* exposed to VOCs from different bacteria. We observed strongest inhibition when LEC was grown on 100% NA and 100% TSA medium, while no substantial VOC activity was observable when LEC was grown on tenfold diluted medium and when grown on SGCA medium (Fig. S3). The antibiotic repertoire used by *L. enzymogenes* has been reported previously to depend on the nutrient abundance and composition in its environment (Lin *et al*., 2021). This is in agreement with other reports, where the total amount of protein (20 g/L for NA and TSA, 3.3 g/L in SGCA) was more important than sugars (no added sugars for NA and TSA, 8.24 g/L for SGCA) in the growth medium (Bruce *et al*., 2003; Lazazzara *et al*., 2017).

Apart from nutrient availability, the volatilome of the VOC emitting microorganism is affected by oxygen availability, pH, and other parameters (Schmidt *et al*., 2015; Lazazzara *et al*., 2017). Oxygen availability might play an important role as several VOCs are metabolic byproducts and reduced oxygen levels can alter metabolic fluxes and overall activity, thereby affecting VOC synthesis (Daft *et al*., 1975; Arras *et al*., 1998; Veselova *et al*., 2019; Tilocca *et al*., 2020). Consistent with this, we observed stronger inhibitory effects under enhanced ventilation (Fig. S5, Table 2). Since *L. enzymogenes* is described as an aerobic organism, higher oxygen supply has been observed to enhance growth and metabolic activity (Lin *et al*., 2021). However, in our experiments optical density was 3- to 6-fold lower under ventilated conditions compared to non-ventilated conditions, indicating that the increase in inhibition was not due to increased bacterial growth. This is in line with observation made on other *Lysobacter* type strains demonstrating that metabolic changes have a much stronger influence on the volatilome than biomass formation (Lazazzara *et al*., 2017). The reduction in cell mass may be related to the lower relative humidity associated with increased air flow, which has previously been reported to affect bacterial growth (Qiu *et al*., 2022). Hence, the increased inhibition under ventilation that we observed might have been caused by enhanced VOC distribution via enhanced air flow and changes in metabolism due to elevated oxygen availability. In summary, the precise mechanisms underlying the observed enhanced growth inhibition under aeration remain to be clarified.

We observed a pH increase of the growth medium from 6.2 to 7.3 due to VOC activity. Vlassi *et al*. (2020) reported an increase in pH from 5.6 to 8.2 upon exposure to VOCs of *L. capsici* AZ78 under similar conditions. The comparable weaker agar alkalization in our experiments might indicate that pH changes play a lesser role in the overall growth inhibition and direct effect of LEC VOCs on the pathogen could be more dominant. However, this hypothesis would still need to be proven.

Besides the inhibition of fungal growth, LEC VOCs lead to intracellular alterations, in particular decreased nuclear abundance and increased lipid accumulation, accompanied by less-defined lipid structures. The presence of multinucleate cells is a well-established characteristic of *R. solani* (Sneh *et al*., 1991) and was observed more frequently in the non-treated cells than in the VOC-treated cells. Potential explanations for reduced nuclear abundance might be either altered nuclear division or stress-induced nuclear degradation (nucleophagy). Cellular self-digestion (autophagy) has been shown for *R. solani* upon treatment with the antifungal volatile agent eugenol (Huang *et al*., 2026). Stress-induced autophagy in connection with nucleophagy has been described in *Aspergillus oryzae* (Shoji *et al*., 2010). The concomitant lipid accumulation strengthens the view of stress-induced processes inside the cell. Lipid droplets are dynamic organelles involved in energy storage and stress adaptation and are known to reorganize under stress conditions and during plant-pathogen interactions (Beccaccioli *et al*., 2019). Lipids have also been described as a means of storage for toxins in fungi upon stress exposure (Chang *et al*., 2015). Furthermore, lipid accumulation has frequently been associated with stress adaptation through ROS-induced stress responses (Shi *et al*., 2017). As VOCs have frequently been described in conjunction with ROS production, this links our observed morphological changes to VOC exposure (Wang *et al*., 2021; Almeida *et al*., 2023).

To identify potential biocontrol-related VOCs, we compared the volatilome of LEC with that of *L. enzymogenes* DSM2043, a closely related strain, but who does not possess biocontrol activity against *R. solani*. We hypothesize that compounds either much higher abundant or exclusively detected in LEC in comparison to DSM2043 contribute to its biocontrol activity against *R. solani*. Previous studies demonstrated the possibility of differentiating bacterial strains, even different *Lysobacter* type strains based on their volatilome fingerprint (Lazazzara *et al*., 2017; Küntzel *et al*., 2018; Rees *et al*., 2018; Beccaria *et al*., 2021). In agreement with this, the volatilomes of LEC and DSM2043 clustered in separate groups in our PCA analysis.

Among the VOCs emitted by both *L. enzymogenes* strains we identified various pyrazines such as dimethylpyrazin, (methyl-)ketones including 6-methyl-2-heptanone and 2-tetradecanone, alcohols such as 2-methyl-1-butanol, the aldehyde benzaldehyde, and alkanes such as decane. Many of them have been widely reported for their antimicrobial effects (Tilocca *et al*., 2020).

As higher or exclusively abundant in LEC we determined 2-undecanone, which has been shown to trigger ROS accumulation in *Sclerotinia sclerotiorum* (Massawe *et al*., 2018). We further identified the derivate undecane, that has been described before to inhibit *P. infestans* growth *in vitro* (Hunziker *et al*., 2015). We also detected 2-heptanone as higher abundant in LEC, which has been shown to inhibit growth of *R. solani in vitro* (Marzouk *et al*., 2021). LEC also produced to a much higher extent 2-decanone, 2-dodecanone and 2-nonanone, that have previously been shown to inhibit mycelial growth of *Fusarium solani in vitro* (Guevara-Avendaño *et al*., 2019). Further 6-meythy-2-heptane induced morphological abnormalities and growth reduction in *A. solani* (Zhang *et al*., 2022), which was also higher abundant in LEC. De Vrieze *et al*. (2015) showed that VOC compounds 2-decanone, 2-undecanone, 2-heptanone produced by *Pseudomonas protegens* were especially effective in inhibiting sporangia germination of *P. infestans in vitro*. While the individual VOCs were not tested separately in this study, their reported biological activities, including against some pathogens we also investigated, provide a basis for interpreting their potential contribution to the observed biocontrol effects.

It has to be noted, that the volatilome we determined might not fully represent the spectrum of the active VOCs causing the observed inhibition against *R. solani*. Ebadzadsahrai *et al*. (2020) found that the volatilome of *C. vaccinii* and the opposing fungus differed depending on whether they were cultivated individually or in co-culture. As many of the identified compounds have already been described in the context of biocontrol, an overlap is highly likely. Definite proof would require co-cultivation, which unfortunately was not feasible here for technical reasons. Further, some of the identified VOCs produced by LEC may have adverse effects depending on their concentration. However, as the GC-MS results are based on relative abundances, no conclusions about potential health risks can be drawn at this stage.

At the same time, the described dynamic nature of the volatilome and the identified influencing factors, such as media composition and oxygen supply provide opportunities for its targeted design and optimization in respect of the use as a biocontrol product. This might enable the enhancement of biocontrol potential while minimizing harmful compounds. Numerous studies have also shown that volatile mixtures can exhibit stronger antifungal activity than individual compounds, with synergistic effects that can substantially enhance the overall inhibition (Kong *et al*., 2022; Shilman *et al*., 2024), a fact that should be considered in biocontrol product development.

Subsequently we observed significant suppression of *P. cubensis* by LEC VOCs on cucumber plants when placing living LEC cultures below the plants. Unfortunately, significant effects were restricted to leaves positioned directly above the VOC source, underlining the need for technical improvements. Increasing VOC abundance did not enhance the protective effect towards neighboring leaves, but instead led to phytotoxicity on the leaves. This needs to be considered when using LEC VOCs for crop protection. Also here, the ability to modulate the VOC composition may provide a means of overcoming this risk.

With regard of potential practical applications, the applicability of VOCs as plant protecting agents has been shown before (Tilocca *et al*., 2020; Belt *et al*., 2026). Calvo *et al*. (2020) demonstrated that VOCs of *Bacillus velezensis* are suitable for post-harvest use in apricots, apples, grapes, and mandarins. In agreement with our data, they observed severe dose-dependent phytotoxicity on the fruits. Furthermore, VOCs can be used as a rhizosphere treatment to control root pathogens, such as *Ralstonia solanacearum* (Raza *et al*., 2016; Belt *et al*., 2026). Tilocca *et al*. (2020) emphasized the use of VOCs especially under air-tight conditions to keep their concentration high and ensure uniform and continuous exposure. This might hint to possible use in greenhouse or foil tunnel crop production. In addition the VOC source could be encapsulated, allowing for a slow and continuous release through micropores of polymer capsules via a diffusion-based release (Jain *et al*., 2024). In combination with applications on low-growing crops and dense canopy structure such as lettuce or potato this might prevent VOCs from dispersing.

The findings of this study further reinforce the versatility of LEC as a biocontrol agent. Furthermore, this study provides new insights into the VOC composition of LEC and their physiological effects on plant pathogens, further advancing our knowledge of their mode of action. However, further research is needed before practical application can be realized, considering factors such as environmental safety, optimization of VOC composition and the identification of ideal exposure conditions.

## Acknowledgements

The authors thank Martin Tröndle for his support with the TEM analysis.

## Data availability statement

The data and the R scripts that were used for figure generation and statistical analysis are available on GitHub at https://github.com/JulianMaierJKI/L.enzymogenes_Volatile_data.

## Author contributions

**J. Maier**: Conceptualization, Data Curation, Formal Analysis, Investigation, Methodology, Software, Visualization, Writing - original draft. **J. Gallinger**: Conceptualization, Formal Analysis, Investigation, Methodology, Writing - review and editing. **M. Molthan**: Investigation, Writing - review and editing. **N. Heil**: Investigation, Methodology, Visualization, Writing – review and editing. **P. Schäfer**: Conceptualization, Resources, Writing - review and editing. **A. Linkies**: Conceptualization, Visualization, Funding Acquisition, Project Administration, Resources, Writing - original draft.

## Supplementary material

Supplementary Material is available online.

