## Supplementary material for "Volatile organic compounds of *L. enzymogenes* LEC suppress the growth of fungal and oomycete pathogens and modify the cell physiology of *Rhizoctonia solani*"

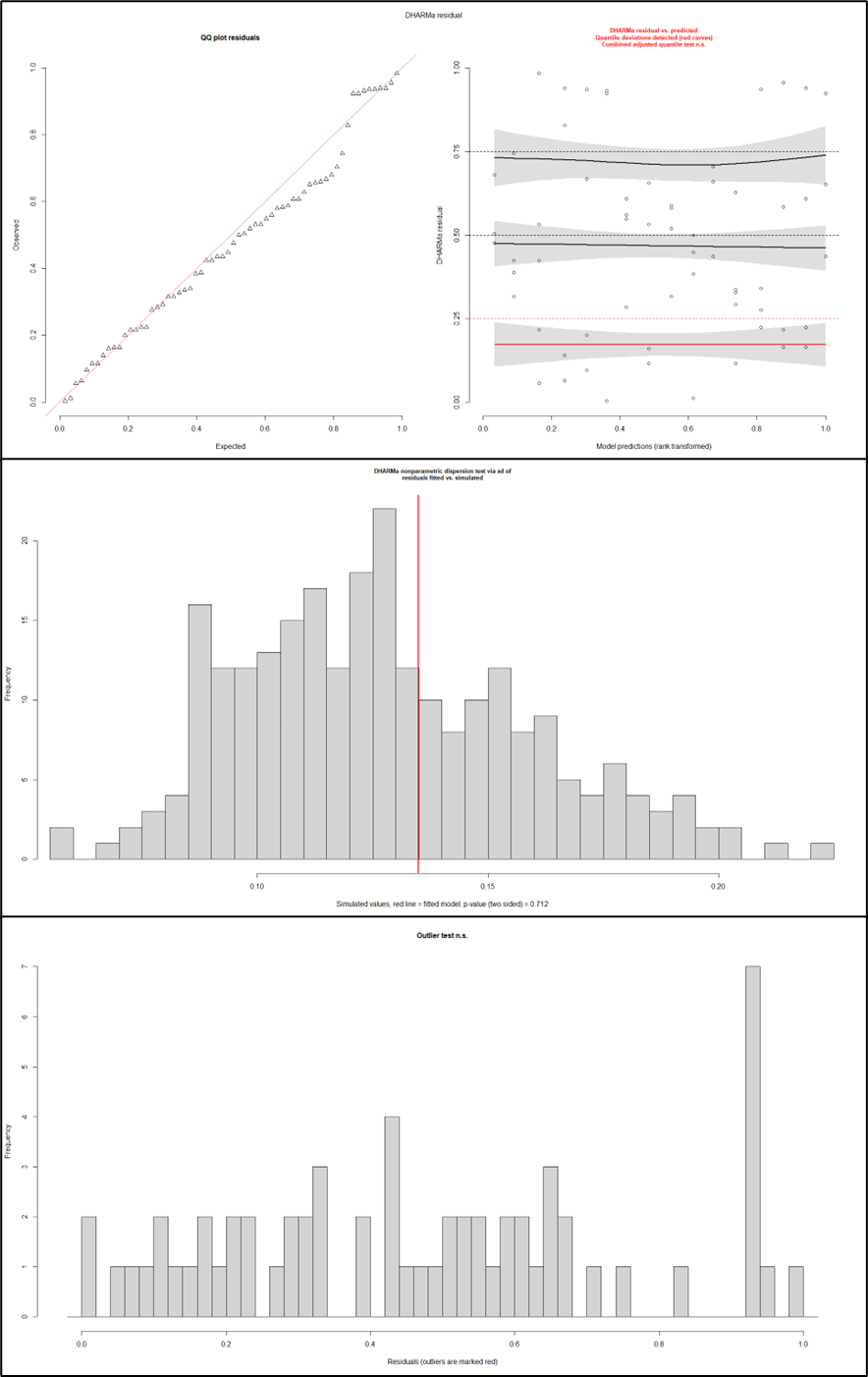

**Fig. S1. Statistical diagnostics of the in vitro experiments. Top)** Shows the output of the DHARMa function plot(simulationOutput). No significant deviation from the assumption of normally distributed residuals is shown. The Kolmogorov–Smirnov test is non-significant. The DHARMa residuals vs. predicted plot shows a slight deviation from the expected values. However, the overall model fit was best for the chosen Gaussian distribution. **Middle)** Shows the output of the DHARMa function testDispersion(). The residuals show no significant skewness and hence no over- or under-dispersion (p-value = 0.712) of the model. **Bottom)** Shows the result of the outliers() test. No significant outliers were detected.

**Table S1. Model specific predicted values and 95% confidence interval of the in vitro experiments.** Shown are the respective predicted absolute growth values in cm based on the chosen model for each pathogen and the corresponding treatment and their 95% confidence interval.

| **Fungi** | **LEC** | **Predicted (cm)** | **(95% CI)** |
| --- | --- | --- | --- |
| *A. radicina* | - | 2.77 | (2.22, 3.31) |
|  | + | 3.61 | (3.56, 3.67) |
| *A. solani* | - | 2.59 | (2.04, 3.13) |
|  | + | 3.64 | (3.58, 3.69) |
| *F. culmorum* | - | 3.61 | (3.06, 4.15) |
|  | + | 3.65 | (3.60, 3.71) |
| *F. graminearum* | - | 3.51 | (2.97, 4.06) |
|  | + | 3.65 | (3.60, 3.70) |
| *P. infestans* | - | 0.18 | (-0.45, 0.81) |
|  | + | 3.66 | (3.61, 3.72) |
| *P. lingam* | - | 0.63 | (0.08, 1.17) |
|  | + | 3.56 | (3.51, 3.62) |
| *P. ultimum* | - | 2.85 | (2.22, 3.48) |
|  | + | 3.61 | (3.56, 3.67) |
| *R. solani* | - | 2.13 | (1.64, 2.62) |
|  | + | 3.63 | (3.57, 3.68) |

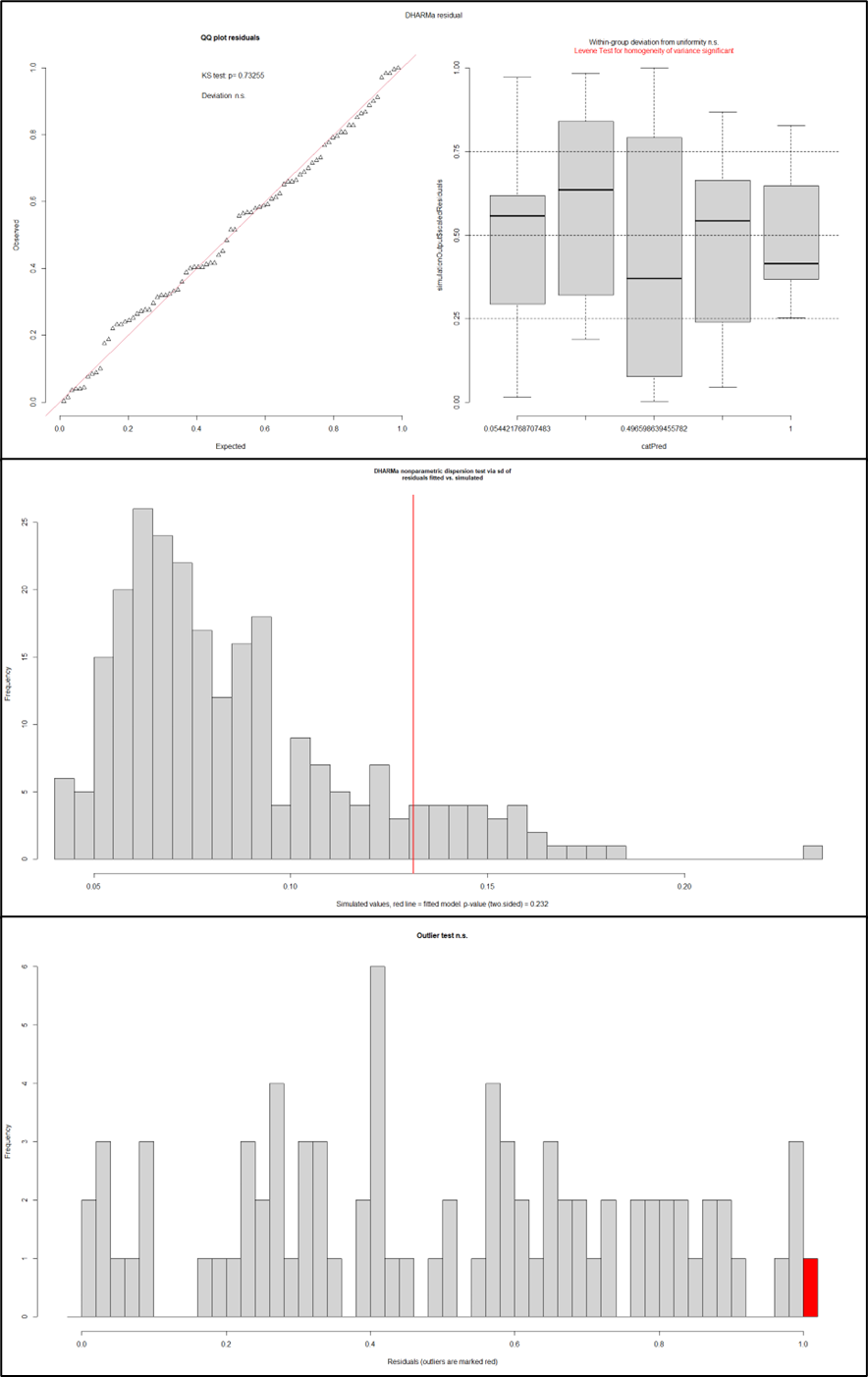

**Fig. S2. Statistical diagnostics of the in planta experiments.** **Top)** Shows the output of the DHARMa function plot(simulationOutput). No significant deviation from the assumption of normally distributed residuals is shown. The Kolmogorov–Smirnov test is non-significant. The Within-group deviation from uniformity is also non-significant. The Levene Test for homogeneity of variance shows a significant deviation and hence the variance differs across groups. However, the overall model fit was best for the chosen Ordbeta distribution. **Middle)** Shows the output of the DHARMa function testDispersion(). The residuals show only a slight skewness but no significant over- or under-dispersion (p-value = 0.264) of the model. **Bottom)** Shows the result of the outliers() test. One significant outlier was detected. However, this outlier represents a true biologically relevant data point and was therefore included in the final model.

**Table S2.** **Model specific predicted values and 95% confidence interval of the in planta experiments.** Shown are the respective predicted values for the relative infected leaf area in % based on the chosen model for each treatment and the corresponding 95% confidence interval.

| **Treatment** | **Predicted (%)** | **(95% CI)** |
| --- | --- | --- |
| CON+ | 94 | (90, 96) |
| CON- | 0 | (0, 3) |
| Cuprozin Progress 0.52% | 8 | (5, 13) |
| - LEC volatiles | 88 | (83, 91) |
| +LEC volatiles | 44 | (37, 51) |

**Table S3. Effect of culture medium of L. enzymogenes LEC on the respective VOC activity, determined as growth inhibition of Alternaria radicina and Rhizoctonia solani.** The trial was done in split-plate Petri dish assays with LEC grown on one side and the fungal isolate on the other side, grown on rye agar. Shown is the radial growth inhibition by LEC VOCs for A. radicina and R. solani in % and the respective culture medium where LEC was propagated on. Each treatment was prepared in duplicates, except for A. radicina-10% NA, A. radicina-10% SGCA and R. solani-10% TSA, which were only tested once. NA: nutrient agar, TSA: tryptone soy agar, SGCA: soybean flour glucose calcium chloride agar.

| **Species** | **LEC growth medium** | **Growth reduction [%]** |
| --- | --- | --- |
| *A. radicina* | 10% NA | 0 |
| *R. solani* | 10% NA | 0 |
| *A. radicina* | 10% SGCA | -0.5 |
| *R. solani* | 10% SGCA | 0 |
| *A. radicina* | 10% TSA | 1 |
| *R. solani* | 10% TSA | 10.8 |
| *A. radicina* | NA | 48.8 |
| *R. solani* | NA | 62.7 |
| *A. radicina* | SGCA | -0.5 |
| *R. solani* | SGCA | 0 |
| *A. radicina* | TSA | 73.9 |
| *R. solani* | TSA | 39.7 |

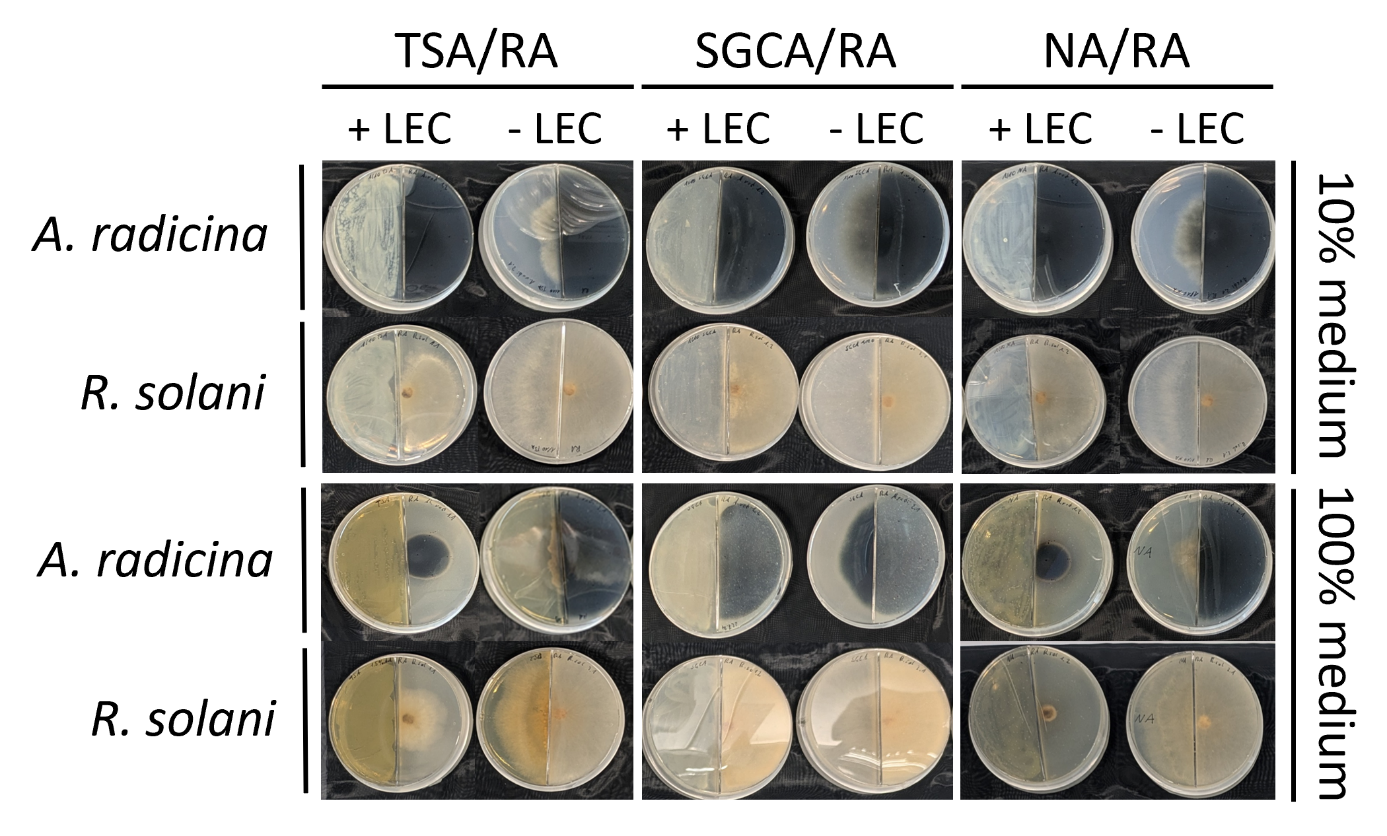

**Fig. S3. Influence of the culture medium on the production of VOCs by Lysobacter enzymogenes isolate LEC.** The inhibitory effect on the radial mycelial growth of Alternaria solani and Rhizoctonia solani is shown. A. solani and R. solani were grown for eight days and four days respectively until the control treatment reached the edge of the plate. LEC: Lysobacter enzymogenes LEC, TSA: tryptic soy agar, NA: nutrient agar, SGCA: soybean flour glucose calcium chloride agar, RA: 10% rye agar. + LEC and - LEC indicate the presence and absence of LEC. Fungal cultures without LEC on the opposing site regularly overgrew the separating barrier of the split-plate.

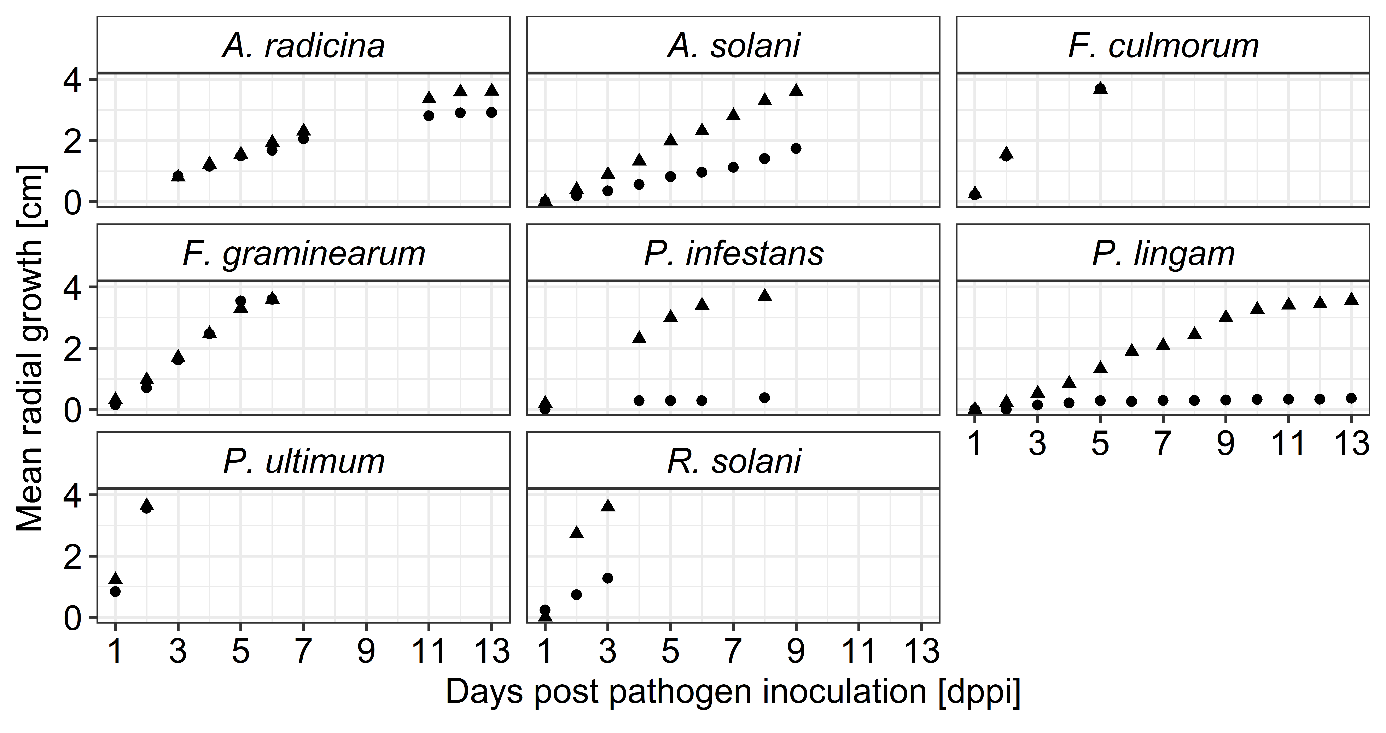

**Fig. S4. Growth of fungal and oomycete isolates over time with and without the exposure of volatile organic compounds (VOCs) of Lysobacter enzymogenes LEC**. Shown is the radial growth (cm) over 13 days for one experimental run, measured daily in three positions per Petri dish with three to four Petri dishes per treatment with (dots) and without (triangles) LEC exposure. A. radicina: Alternaria radicina, A. solani: Alternaria solani, F. culmorum: Fusarium culmorum, F. graminearum: Fusarium graminearum, P. infestans: Phytophthora infestans, P. lingam: Phoma lingam, P. ultimum: Pythium ultimum, R. solani: Rhizoctonia solani.

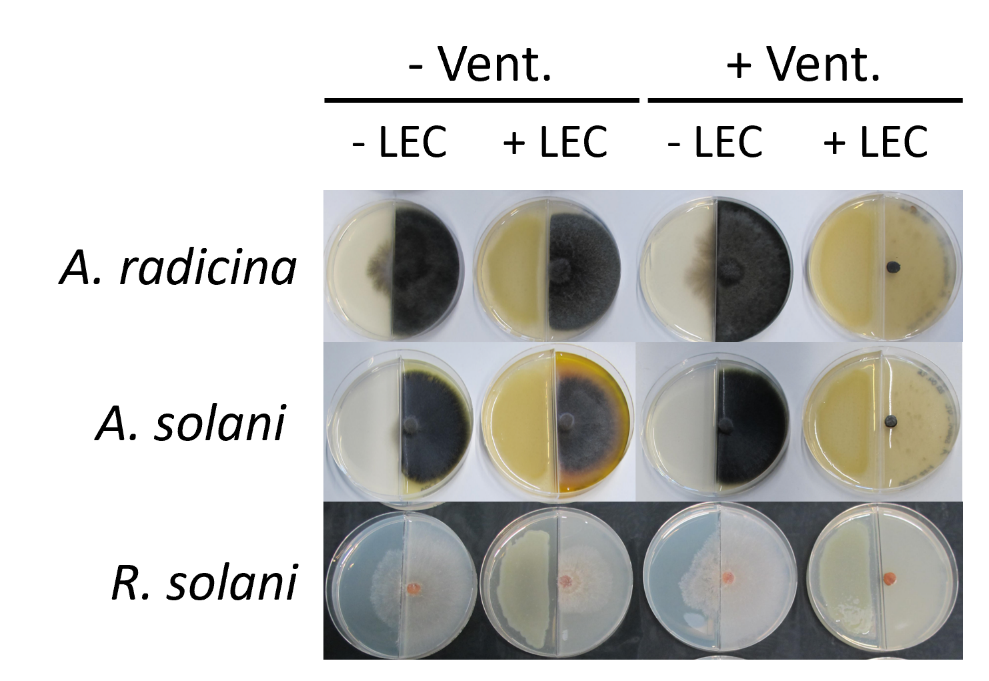

**Fig. S5. Influence of increased ventilation conditions on the radial growth reduction of three phytopathogenic fungi caused by volatile organic compounds (VOCs) of Lysobacter enzymogenes LEC.** The inhibitory effect on the radial mycelial growth of Alternaria radicina, Alternaria solani and Rhizoctonia solani is shown. The pictures show the fungi after nine days (A. radicina and A. solani), 10 days (A. solani, – Vent.) and 3 days (R. solani) of growth. LEC: Lysobacter enzymogenes isolate LEC. + LEC and - LEC indicate the presence and absence of LEC. – Vent. and + Vent. indicate increased ventilation conditions. Fungi grown under control conditions regularly crossed the separating barrier of the split-plate growing onto the nutrient agar side (- LEC).

**Table S4. VOC-dependent changes of medium pH.** pH values of the rye agar surface were measured in split-plate assays with (LEC) and without (Ctrl) the exposure to L. enzymogenes LEC three days after LEC inoculation. Values from three independent experiments are shown, together with mean values (Mean) and standard deviation (SD).

| **Experiment** | **Ctrl** |  | **LEC** |  |
| --- | --- | --- | --- | --- |
| 1 | 6.0 |  | 7.5 |  |
| 2 | 6.0 |  | 7.0 |  |
| 3 | 6.5 | **SD** | 7.3 | **SD** |
| **Mean** | 6.2 | 0.29 | 7.3 | 0.25 |

**Table S5. Screening of Lysobacter isolates for the identification of a GC-MS control.** L. enzymogenes (LEC and DSM2043), L. antibioticus (DSM 2044) and L. gummosus (DSM 6980) were grown on NA and their suppressive volatile activity was analysed in split-plate assays against Rhizoctonia solani (R. solani), Phytophthora infestans (P. infestans) and Phoma lingam (P. lingam) gown on 10% rye agar. Shown is the mean radial growth reduction (% and the number of replicate split-plates (n).

| **Fungi** | **Bacteria** | **Isolate** | **Growth reduction [%]** | **n** |
| --- | --- | --- | --- | --- |
| *R. solani* | *L. enzymogenes* | DSM 2043 | 0 | 3 |
| *R. solani* | *L. antibioticus* | DSM 2044 | 0 | 3 |
| *R. solani* | *L. gummosus* | DSM 6980 | 0 | 3 |
| *P. infestans* | *L. enzymogenes* | DSM 2043 | 92 | 3 |
| *P. infestans* | *L. antibioticus* | DSM 2044 | 89 | 3 |
| *P. infestans* | *L. gummosus* | DSM 6980 | 89 | 2 |
| *P. lingam* | *L. enzymogenes* | DSM 2043 | 71 | 3 |
| *P. lingam* | *L. antibioticus* | DSM 2044 | 74 | 2 |
| *P. lingam* | *L. gummosus* | DSM 6980 | 71 | 2 |

**Table S6. Results of the statistical analysis of the GC-MS analysis.** Left) Table of the Kruskal-Wallis-Test results showing the identified volatile organic compounds (VOC) and the respective p-value (p-value) and the Benjamini-Hochberg adjusted p-values (P.adj). Right) Table of the Dunn-Test results showing the identified volatile organic compounds (VOC), the respective comparison between bacterial organisms and the media control (C: Control medium (Nutrient Agar), D: DSM2043, L: LEC), the respective Z-value (Z) and the Benjamini-Hochberg adjusted p-values (P.adj).

| **Kruskal-Wallis Results** | | | **Dunn-Test Results** | | | |
| --- | --- | --- | --- | --- | --- | --- |
| **VOC** | **p-value** | **P.adj** | **VOC** | **Comparison** | **Z** | **P.adj** |
| 2-ethylhexanol | 0.463 | 0.653 | 2-decanone | C - L | -3.654 | 0.000 |
| 2,4,4-trimethylpentene | 0.697 | 0.822 | 2-decanone | D - L | -3.963 | 0.000 |
| 2-decanone | 0.000 | 0.000 | 2-heptanone | C - L | -3.707 | 0.000 |
| 2-heptanone | 0.000 | 0.000 | 2-heptanone | D - L | -3.727 | 0.001 |
| 6-methyl-2-heptanone | 0.000 | 0.000 | 6-methyl-2-heptanone | C - L | -3.774 | 0.000 |
| 2-methyl-1-butanol | 0.001 | 0.003 | 6-methyl-2-heptanone | D - L | -3.520 | 0.001 |
| 2-nonanone | 0.000 | 0.000 | 2-methyl-1-butanol | C - D | -3.550 | 0.001 |
| RI 1563 | 0.028 | 0.054 | 2-methyl-1-butanol | C - L | -2.845 | 0.007 |
| 3-furaldehyde | 0.000 | 0.000 | 2-nonanone | C - L | -3.396 | 0.001 |
| acetophenone | 0.078 | 0.126 | 2-nonanone | D - L | -3.827 | 0.000 |
| benzaldehyde | 0.000 | 0.001 | 3-furaldehyde | C - D | 4.064 | 0.000 |
| cyclopentanone | 0.883 | 0.883 | 3-furaldehyde | C - L | 4.164 | 0.000 |
| decane | 0.010 | 0.020 | benzaldehyde | C - D | 3.372 | 0.001 |
| 2-methoxyfuran | 0.811 | 0.883 | benzaldehyde | C - L | 3.830 | 0.000 |
| n-dodecane | 0.000 | 0.001 | decane | C - D | 2.681 | 0.011 |
| p-xylene | 0.480 | 0.653 | decane | C - L | 2.747 | 0.018 |
| phenylacetaldehyde | 0.000 | 0.000 | n-dodecane | C - D | 4.077 | 0.000 |
| pyrazin (RI 919) | 0.045 | 0.076 | n-dodecane | C - L | 2.627 | 0.013 |
| pyrazin (RI 1052) | 0.003 | 0.007 | phenylacetaldehyde | C - D | 4.064 | 0.000 |
| pyrazin (RI 1172) | 0.156 | 0.242 | phenylacetaldehyde | C - L | 4.164 | 0.000 |
| pyrazin (RI 1058) | 0.000 | 0.000 | pyrazin (RI 1052) | C - D | -2.769 | 0.008 |
| RI 1116 | 0.866 | 0.883 | pyrazin (RI 1052) | C - L | -3.278 | 0.003 |
| RI 1170 | 0.502 | 0.656 | pyrazin (RI 1058) | C - L | -4.038 | 0.000 |
| RI 1232 | 0.037 | 0.066 | pyrazin (RI 1058) | D - L | -3.011 | 0.004 |
| undecanone | 0.000 | 0.000 | undecanone | C - L | -3.346 | 0.001 |
| 2-dodecanone | 0.000 | 0.001 | undecanone | D - L | -4.021 | 0.000 |
| RI 1401 | 0.002 | 0.004 | 2-dodecanone | C - L | -3.525 | 0.001 |
| RI 1473 | 0.028 | 0.054 | 2-dodecanone | D - L | -3.328 | 0.001 |
| RI 1601 | 0.005 | 0.012 | RI 1401 | C - D | 3.167 | 0.002 |
| RI 875 | 0.613 | 0.772 | RI 1401 | C - L | 3.245 | 0.004 |
| RI 884 | 0.701 | 0.822 | RI 1601 | C - D | 2.576 | 0.015 |
| RI 931 | 0.838 | 0.883 | RI 1601 | C - L | 3.100 | 0.006 |
| RI 949 | 0.755 | 0.856 |  |  |  |  |
| xylene | 0.231 | 0.341 |  |  |  |  |

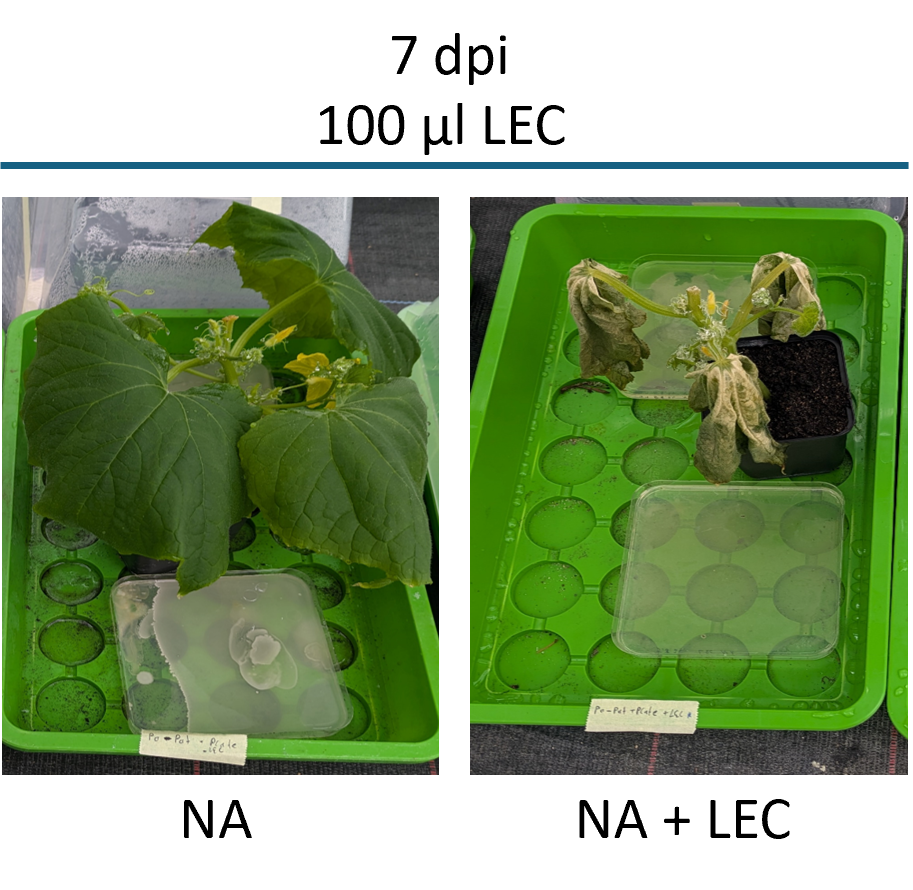

**Fig. S6. Phytotoxicity of Lysobacter enzymogenes LEC volatile organic compounds on cucumber plants.** Petri dishes spot-inoculated with 100 µl of a LEC suspension (NA + LEC; 10^7 cells/ml) or empty media (NA, 0.6% NaCl) were placed in closed propagation boxes with four-weeks old cucumber plants. Each treatment was done in duplicate, pictures of representative plants are shown seven days after placement of the LEC culture. The control agar plate shows growth of fungi or bacteria due to water and air exposure during the experiment. The LEC plate appears empty due to the almost clear growth phenotype of LEC on agar plates.
